# The amyloid precursor protein regulates synaptic transmission at medial perforant path synapses

**DOI:** 10.1101/2022.09.05.506635

**Authors:** Maximilian Lenz, Amelie Eichler, Pia Kruse, Christos Galanis, Dimitrios Kleidonas, Peter Jedlicka, Ulrike Müller, Thomas Deller, Andreas Vlachos

## Abstract

The perforant path provides the main cortical excitatory input to the hippocampus. Due to its important role in information processing and coding, entorhinal projections to the dentate gyrus have been studied in considerable detail. Nevertheless, a characterization of synaptic transmission between individual connected pairs of entorhinal stellate cells and dentate granule cells is still pending. Here, we have used organotypic entorhino-hippocampal tissue cultures, in which the entorhino-dentate (EC-GC) projection is present and EC-GC pairs can be studied using whole-cell patch clamp recordings. Using cultures of wildtype mice, the properties of EC-GC synapses formed by afferents from the lateral and medial entorhinal cortex were compared and differences in short-term plasticity were revealed. Since the perforant path is severely affected in Alzheimer’s disease, we used cultures of APP-deficient mice to address the role of the amyloid-precursor protein (APP) at this synapse. APP-deficiency caused alterations in excitatory neurotransmission at medial perforant path synapses that were accompanied by transcriptomic and ultrastructural changes. Moreover, the deletion of pre- but not postsynaptic APP through the local injection of Cre-expressing AAVs in conditional APP^flox/flox^ tissue cultures increased the efficacy of neurotransmission at perforant path synapses. Together, these data suggest a physiological role for presynaptic APP at medial perforant path synapses, which may be adversely affected under conditions of altered APP processing.

## INTRODUCTION

The dentate gyrus is considered the gateway to the hippocampus (Hsu, 2007; Krook-Magnuson et al., 2015; Winson and Abzug, 1977), receiving input from cortical areas and recurrent fibers from hippocampal regions (Amaral et al., 2007; Patton and McNaughton, 1995; Ribak and Shapiro, 2007). Signal integration and processing in this region is thus of considerable interest in understanding dedicated hippocampal features (Jonas and Lisman, 2014; Leutgeb et al., 2007; Leutgeb and Moser, 2007), e.g. memory formation (Trimper et al., 2017), orientation in space and time (GoodSmith et al., 2017) and implementation of complex behavior (Hainmueller and Bartos, 2018; Senzai and Buzsaki, 2017).

Dentate granule cells, the predominant cell type within the dentate gyrus, receive their major excitatory inputs from either the perforant path or recurrent fibers that originate from the hilar region (Amaral et al., 2007). The perforant path originating from layer 2 principal neurons of the entorhinal cortex forms asymmetric synapses at granule cell dendrites in the outer two thirds of the molecular layer (Forster et al., 2006). These axons reach the entire transverse extent of the molecular layer, while their projection along the septotemporal axis remains limited (Tamamaki and Nojyo, 1993). In multiple studies, distinct features of this projection, such as short-term plasticity expression have been identified using, e.g., local electric stimulation and field potential recordings (Jedlicka et al., 2012; Petersen et al., 2013; Winkels et al., 2009). However, an investigation of perforant path synapses on the level of individual pairs of neurons, i.e., layer 2 stellate cells in the entorhinal cortex and dentate granule cells, is lacking.

Beyond its role in normal brain function, the perforant path has been recognized as a major target for memory formation and neuropsychiatric diseases (Kirkby and Higgins, 1998; Roy et al., 2016; Smith and McMahon, 2018). Among these diseases, Alzheimer’s disease is associated with progressive alterations along the perforant path, which correlates with cognitive decline (Robinson et al., 2014). In this context, the processing of the amyloid precursor protein (APP) and specifically “synaptotoxic” amyloid-β (Aβ) has been linked to pathognomonic plaque formation and cognitive decline in Alzheimer’s disease. Moreover, proteolytic amyloidogenic APP cleavage at perforant path terminals was found to act as a regulator of synaptic transmission and hippocampal function (Harris et al., 2010; Lazarov et al., 2002; Soldano and Hassan, 2014). Nevertheless, the impact of APP on synaptic transmission at individual perforant path synapses remains elusive.

Here, we addressed the physiological role of APP for synaptic transmission at the EC-GC synapse. Using paired whole-cell patch-clamp recordings in wildtype and APP-deficient organotypic entorhino-hippocampal tissue cultures, we demonstrate the distinct electrophysiological features of excitatory inputs to dentate granule cells – including the hilar mossy cell pathway. We provide evidence that presynaptic (but not postsynaptic) APP-deficiency is accompanied by an increase in the efficacy of excitatory synaptic transmission between connected pairs of neurons at the medial perforant path. Consistent with these findings, ultrastructural analysis revealed a higher number of docked vesicles at presynaptic active zones in the molecular layer of the APP-deficient dentate gyrus. These results disclose the synaptic properties of the entorhino-hippocampal pathway at the level of individual connected pairs of neurons and demonstrate that presynaptic APP is a key regulator of excitatory synaptic transmission at perforant path synapses in the dentate gyrus.

## MATERIALS AND METHODS

### Ethics statement

Mice were maintained in a 12 hour light/dark cycle with food and water available *ad libitum*. Every effort was made to minimize distress and pain of animals. All experimental procedures were performed according to the German animal welfare legislation and approved by the animal welfare committee and/or the animal welfare officer at the Goethe-University Frankfurt, Faculty of Medicine, and the Albert-Ludwigs-University Freiburg, Faculty of Medicine.

### Preparation of organotypic tissue cultures

Entorhino-hippocampal tissue cultures were prepared at postnatal day 4 - 5 from C57BL/6J and APP-deficient (APP^−/−^; (Li et al., 1996)) animals as previously described (Del Turco and Deller, 2007). Conditional APP-deficient cultures were prepared from homozygous APP^flox/flox^ animals (Mallm et al., 2010) that were crossed to B6.Cg-Gt(ROSA)26Sor^tm14(CAG-tdTomato)Hze^/J (Ai14^+/−^; Jackson Laboratories #007914; (Madisen et al., 2010)). The newly generated mouse line (Ai14-APP^flox/flox^) is a reporter mouse revealing nuclear recombination by cellular tdTomato expression. Cultivation medium contained 50 % (v/v) MEM, 25 % (v/v) basal medium eagle, 25 % (v/v) heat-inactivated normal horse serum, 25 mM HEPES buffer solution, 0.15 % (w/v) bicarbonate, 0.65 % (w/v) glucose, 0.1 mg ml^−1^ streptomycin, 100 U ml^−1^ penicillin, and 2 mM glutamax. The pH was adjusted to 7.3 and the medium was replaced three times per week. All tissue cultures were allowed to mature for at least 18 days in humidified atmosphere with 5 % CO_2_ at 35 °C, since at this age a steady state in structural and functional properties of the organotypic tissue cultures is reached (Hailer et al., 1996; Humpel, 2015; Vlachos et al., 2012a; Vlachos et al., 2013).

### Perforant path tracing and local viral Cre-GFP expression

Adeno-associated viruses (AAV) obtained from SignaGen Laboratories, Maryland (AAV2-Synapsin-tdTOMATO, #SL100896 and AAV2-Synapsin-GFP, #SL100817) and the University of North Carolina Vector Core (UNC Vector Core; AAV2-hSyn-GFP-Cre) were injected into the entorhinal cortex at 3 - 5 days *in vitro* at a Zeiss Axioscope 2 equipped with a 4x objective (air, NA 0.1) using borosilicate glass pipettes (c.f. (Lenz et al., 2019)). Cultures were returned to the incubator immediately after injection and allowed to mature for at least 18 days in a humidified atmosphere with 5 % CO_2_ at 35 °C.

### Immunohistochemistry

Cultures were fixed in a solution of 4 % (w/v) paraformaldehyde (PFA) in phosphate-buffered saline (PBS, 0.1 M, pH 7.4) and 4 % (w/v) sucrose for 1 h. Fixed cultures were incubated for 1 h with 10 % (v/v) normal goat serum (NGS) in 0.5 % (v/v) Triton X-100-containing PBS to block non-specific staining. To label calretinin, whole tissue cultures were incubated with rabbit anti calretinin (1:1000; Synaptic Systems, #214102) in PBS containing 10 % (v/v) normal goat serum (NGS) and 0.1 % (v/v) Triton X-100 at 4°C overnight. Cultures were washed and incubated for 3 h with appropriate secondary antibodies (1:1000, in PBS with 10 % NGS or NHS, 0.1 % Triton X-100; Invitrogen). TO-PRO^®^ or DAPI (1:5000 in PBS for 10 min; TO-PRO^®^: Invitrogen, #T-3605; DAPI: Thermo Scientific, #62248) nuclear stain was used to visualize cytoarchitecture. Sections were washed, transferred onto glass slides and mounted for visualization with anti-fading mounting medium (DAKO Fluoromount).

Confocal images in immunostainings were acquired using a Leica SP8 confocal microscope equipped with a 60x objective lens (NA 1.4, Leica).

### Posthoc-staining

Cultures were fixed in a solution of 4 % (w/v) paraformaldehyde (PFA) in phosphate-buffered saline (PBS, 0.1 M, pH 7.4) and 4 % (w/v) sucrose for 1 h. Fixed cultures were incubated for 1 h with 10 % (v/v) normal goat serum (NGS) in 0.5 % (v/v) Triton X-100-containing PBS. Biocytin filled cells were counterstained with Alexa 488- or Alexa 647-conjugated streptavidin (1:1000 in PBS with 10 % NGS, 0.1 % Triton X-100; Invitrogen, #S-32354 and #S-32357 respectively) for 4 h and DAPI or TO-PRO^®^ staining was used to visualize cytoarchitecture (1:5000 in PBS for 10 min; TO-PRO^®^: Invitrogen, #T-3605; DAPI: Thermo Scientific, #62248). Slices were washed, transferred, and mounted onto glass slides for visualization with anti-fading mounting medium (DAKO Fluoromount). Confocal images were acquired using a Nikon Eclipse C1si laser-scanning microscope with a 4x objective lens (NA 0.2, Nikon), a 20x objective lens (NA 0.9, Nikon) and a 60x objective lens (NA 1.4, Nikon) or a Leica SP8 confocal microscope equipped with a 40x objective lens (NA 1.3, Leica). Detector gain and amplifier were initially set to obtain pixel intensities within a linear range.

### Transmission Electron Microscopy

APP^+/+^ and APP^−/−^ tissue cultures were fixed in 4 % paraformaldehyde (w/v) and 2 % glutaraldehyde (w/v) in 0.1 M phosphate buffer (PB) overnight and washed for 1 hour in 0.1 M PB. After fixation, tissue cultures were sliced with a vibratome and the slices were incubated with 1 % osmium tetroxide for 20 min in 5 % (w/v) sucrose containing 0.1 M PB. The slices were washed 5 times for 10 min in 0.1 M PB and washed in graded ethanol (10 min in 10 % (v/v) and 10 min in 20 (v/v)). The slices were then incubated with uranyl acetate (1 % (w/v) in 70 % (v/v) ethanol)) overnight and subsequently dehydrated in graded ethanol 80 % (v/v), 90 % (v/v) and 98 % (v/v) for 10 min. Finally, slices were incubated with 100 % (v/v) ethanol two times for 15 min followed by two 15 min washes with propylene oxide. The slices were then transferred for 30 min in a 1:1 mixture of propylene oxide with durcupan and then for 1 hour in durcupan. The durcupan was exchanged for fresh durcupan and the slices were transferred in 4 °C overnight. The slices were then embedded between liquid release-coated slides and coverslips. Cultures were re-embedded in blocks and ultrathin sections were collected on copper grids. Electron microscopy was performed with a LEO 906E microscope (Zeiss) at 4646x magnification. Acquired images were saved as TIF-files and analyzed using the ImageSP Viewer software (http://e.informer.com/sys-prog.com). In each group, 50 synapses from 5 independent tissue cultures were analyzed in the distal parts of the molecular layer. Asymmetric spine synapses were identified and the total amount of presynaptic vesicles and docked vesicles to presynaptic active zones was manually quantified by an investigator blind to the genotype.

### Paired whole-cell patch-clamp recordings

Whole-cell voltage-clamp recordings from dentate granule cells of slice cultures were carried out at 35 °C (2-5 neurons per culture). The bath solution contained 126 mM NaCl, 2.5 mM KCl, 26 mM NaHCO_3_, 1.25 mM NaH_2_PO_4_, 2 mM CaCl_2_, 2 mM MgCl_2_, and 10 mM glucose. For EPSC recordings patch pipettes contained 126 mM K-gluconate, 4 mM KCl, 4 mM Mg-ATP, 0.3 mM Na_2_-GTP, 10 mM PO-creatine, 10 mM HEPES and 0.3 % (w/v) biocytin (pH = 7.25 with KOH, 290 mOsm with sucrose) having a tip resistance of 4-6 MΩ. In some experiments Alexa488 was added to the internal solution to visualize neuronal morphology prior to recordings. Cells were visually identified using an LN-Scope (Luigs and Neumann, Ratingen, 5 Germany) or a Zeiss LSM750 equipped with infrared dot-contrast and a 40× water-immersion objective (numerical aperture [NA] 0.8; Olympus). Electrophysiological signals were amplified using a Multiclamp 700B amplifier, digitized with a Digidata 1550B digitizer, and visualized with the pClamp 11 software package. Neurons were recorded at a holding potential of −70 mV. Recordings were discarded if leak current or series resistance changed significantly and/or reached ≥ 30 MΩ. Action potentials were generated in the presynaptic cell by 5 ms square current pulses (1 nA) elicited at 0.2 Hz (up to 50 pulses), while recording unitary excitatory postsynaptic currents (uEPSCs) from dentate granule cells. Neurons were considered to be connected if > 5 % of action potentials evoked time-locked inward uEPSCs. For short-term plasticity, 5 action potentials were applied at 10 or 20 Hz, respectively (inter-sweep-interval: 5 sec, 30 repetitions).

### Regional mRNA library preparation and transcriptome analysis

RNA library preparations for transcriptome analysis were performed using the NEBNext® Single Cell/Low Input RNA Library Prep Kit for Illumina^®^ (New England Biolabs, #E6420) according to the manufacturer’s instructions. Briefly, isolation of the dentate gyrus from individual tissue cultures was performed using a scalpel. One isolated dentate gyrus was transferred to 7.5 μl lysis buffer (supplemented with murine RNase inhibitor) and homogenized using a pestill. Samples were centrifuged for 30 seconds at 10000g and 5 μl of supernatant were collected from individual samples and further processed. After cDNA synthesis, cDNA amplification was performed according to the manufacturer’s protocol with 12 PCR cycles. The cDNA yield was subsequently analyzed by a High Sensitivity DNA assay on a Bioanalyzer instrument (Agilent). The amount of cDNA was adjusted to 10 ng for further downstream applications. After fragmentation and adaptor ligation, dual index primers (New England Biolabs, #E7600S) were ligated in a library amplification step using 10 PCR cycles. Libraries were finally cleaned up with 0.8X SPRI beads following a standard bead purification protocol. Library purity and size distribution were assessed with a High Sensitivity DNA assay on a Bioanalyzer instrument (Agilent). We quantified the libraries using the NEBNext Library Quant Kit for Illumina (New England Biolabs, #E7630) based on the mean insert size provided by the Bioanalyzer. A 10 nM sequencing pool (120 μl in Tris-HCl, pH 8.5) was generated for sequencing on the NovaSeq6000 sequencing platform (Illumina; service provided by CeGaT GmbH, Tübingen, Germany). We performed a paired-end sequencing with 151 bp read length. Data analysis was performed at the Galaxy platform (usegalaxy.eu; (Galaxy, 2022)). All files contained more than 10 M high-quality reads (after mapping to the reference genome; mm10) with a phred quality of at least 30 (>90% of total reads).

### Compartmental modeling

Compartmental simulations of synaptic EPSCs in dentate granule cells were performed as described before (Vlachos et al., 2012b) using the simulation environment NEURON ((Hines and Carnevale, 1997); www.neuron.yale.edu). Briefly, we used 8 reconstructed morphologies of mouse dentate granule cells (Schmidt-Hieber et al., 2007) from ModelDB (accession number 95960). Passive biophysical properties were implemented based on data and modeling of Schmidt-Hieber and colleagues (Schmidt-Hieber et al., 2007). Excitatory (AMPA-like) synaptic currents were simulated using a double-exponential conductance change with a peak amplitude of 0.5 nS, a rise time of 0.2 ms, a decay time of 2.5 ms, a peak conductance of 0.5 nS and a reversal potential of 0 mV. To determine the dependence of simulated EPSC amplitudes on the distance from the soma, identical single synaptic input was activated at 6 different locations along a path between the soma and a distal end of the dendrite, and corresponding EPSCs were detected at the soma. Simulated cells were voltage clamped at −70 mV. To simulate compound EPSCs, we monitored voltage-clamped somatic currents in 8 model granule cells in which synaptic activity was triggered by synchronous activation of dendritic AMPA synapses placed at equidistant locations in all dendritic branches.

### Quantification and statistics

Electrophysiological data were analyzed using pClamp 10.7 (Axon Instruments) and MiniAnalysis (Synaptosoft) software. The fraction of action potentials not followed by time-locked excitatory postsynaptic current responses was considered as synaptic failure rate. The uEPSC amplitude, risetime (rise50), and area were assessed in uEPSCs from successfully transmitted action potentials, as well as the mean amplitude of all successfully evoked postsynaptic responses. For evaluation of short-term-plasticity experiments, recorded traces were averaged in Clampfit 10.7, irrespective of successful synaptic transmission at individual pulses. In some analyses, postsynaptic responses were normalized to the first response in averaged traces.

Presynaptic ultrastructural features were analyzed in randomly selected perforant path terminals from electron micrographs of the outer parts of the molecular layer. Postsynaptic features of the same set of images were analyzed in a previous study (Galanis et al., 2021). Presynaptic terminals and vesicles were manually assessed by an investigator blind to experimental conditions.

RNA sequencing data were uploaded to the galaxy web platform (public server: usegalaxy.eu; Afgan et al., 2018; Jalili et al., 2020; Afgan et al., 2016) and transcriptome analysis was performed using the Galaxy platform in accordance with the reference-based RNA-seq data analysis tutorial (Batut et al., 2021). Adapter sequences, low quality, and short reads were removed via the CUTADAPT tool (Galaxy version 3.5+galaxy0). Reads were mapped using RNA STAR (Galaxy version 2.7.8a+galaxy0) with the mm10 Full reference genome (Mus musculus). The evidence-based annotation of the mouse genome (GRCm38), version M25 (Ensembl 100) served as gene model (GENCODE). For an initial assessment of gene expression, unstranded FEATURECOUNT (Galaxy version 2.0.1+galaxy2) analysis was performed from RNA STAR output. Only samples that contained >60% uniquely mapping reads (feature: “exon”) were considered for further analysis. Statistical evaluation was performed using DESeq2 (Galaxy version 2.11.40.7+ galaxy1) with “genotype” as the primary factor that might affect gene expression. Genes with a low number of mean reads (< 150 counts) were excluded from further analysis. Genes were considered as differentially expressed if the adjusted p-value was < 0.05. Data visualization was performed according to a modified version of a previously published workflow (Reimand et al., 2019). A ranked list for the gene set enrichment analysis was generated based on Wald-Stats from the DeSeq2 output. For the consecutive gene set enrichment analysis, we used the GSEA software package (BROAD institute; version GSEA_4.2.1) with integrated MSigDB gene sets. Enrichment maps were built using Cytoscape (version 3.9.0) with the EnrichmentMap application. Heatmaps were generated based on z-scores of the normalized count table.

Data were statistically analyzed using GraphPad Prism 7 (GraphPad software, USA). For statistical comparison of two experimental groups, a Mann-Whitney-test was employed. For the evaluation of data sets with three experimental groups, a Kruskal-Wallis test followed by Dunn’s posthoc correction was performed. Short-term plasticity experiments were statistically assessed by the repeated measure (RM) two-way ANOVA test with Sidak’s (two groups) or Tukey’s (three groups) multiple comparisons test. uEPSC amplitude values from individual cells were stacked in subcolumns and pulse number defined tabular rows (COLUMN factor: pathway, genetic background or recombination; ROW factor: pulse number). p-values < 0.05 were considered statistically significant (*p < 0.05, **p < 0.01, ***p < 0.001). Results that did not yield significant differences were designated ‘ns’. Statistical differences in XY-plots were indicated in the legend of the figure panels (*) when detected through multiple comparisons, irrespective of their localization and their level of significance. In the text and figures, values represent the mean ± standard error of the mean (s.e.m.).

### Digital illustrations

Confocal image stacks were stored as TIF files. Figures were prepared using the ImageJ software package (https://imagej.nih.gov/ij/) and Photoshop graphics software (Adobe, San Jose, CA, USA). Image brightness and contrast were adjusted.

## RESULTS

### Laminar innervation of the dentate gyrus in organotypic entorhino-hippocampal tissue cultures

The perforant path was visualized in the dentate gyrus of organotypic entorhino-hippocampal tissue cultures using local viral injections of adeno-associated viral vectors that express either GFP or tdTomato under the control of a synapsin promotor. Injections were performed in the medial (tdTomato-expression) and the lateral part (GFP-expression) of the entorhinal cortex in the same set of cultures (Figure 1A). Indeed, GFP-positive axons originating from neurons in the lateral part of the cultured entorhinal cortex were readily detected in the outer molecular layer (oml) of the dentate gyrus, while in the middle molecular layer (mml) in the same set of cultures tdTomato-labeled axons were observed (Figure 1B). In addition, mossy cell fibers were visualized in the inner molecular layer (iml) using calretinin immunostainings in AAV-traced tissue cultures (Figure 1B), confirming the organotypic, i.e., *in vivo-*like, laminar innervation of the dentate gyrus (Forster et al., 2006; Frotscher and Heimrich, 1995).

**Figure 1:**
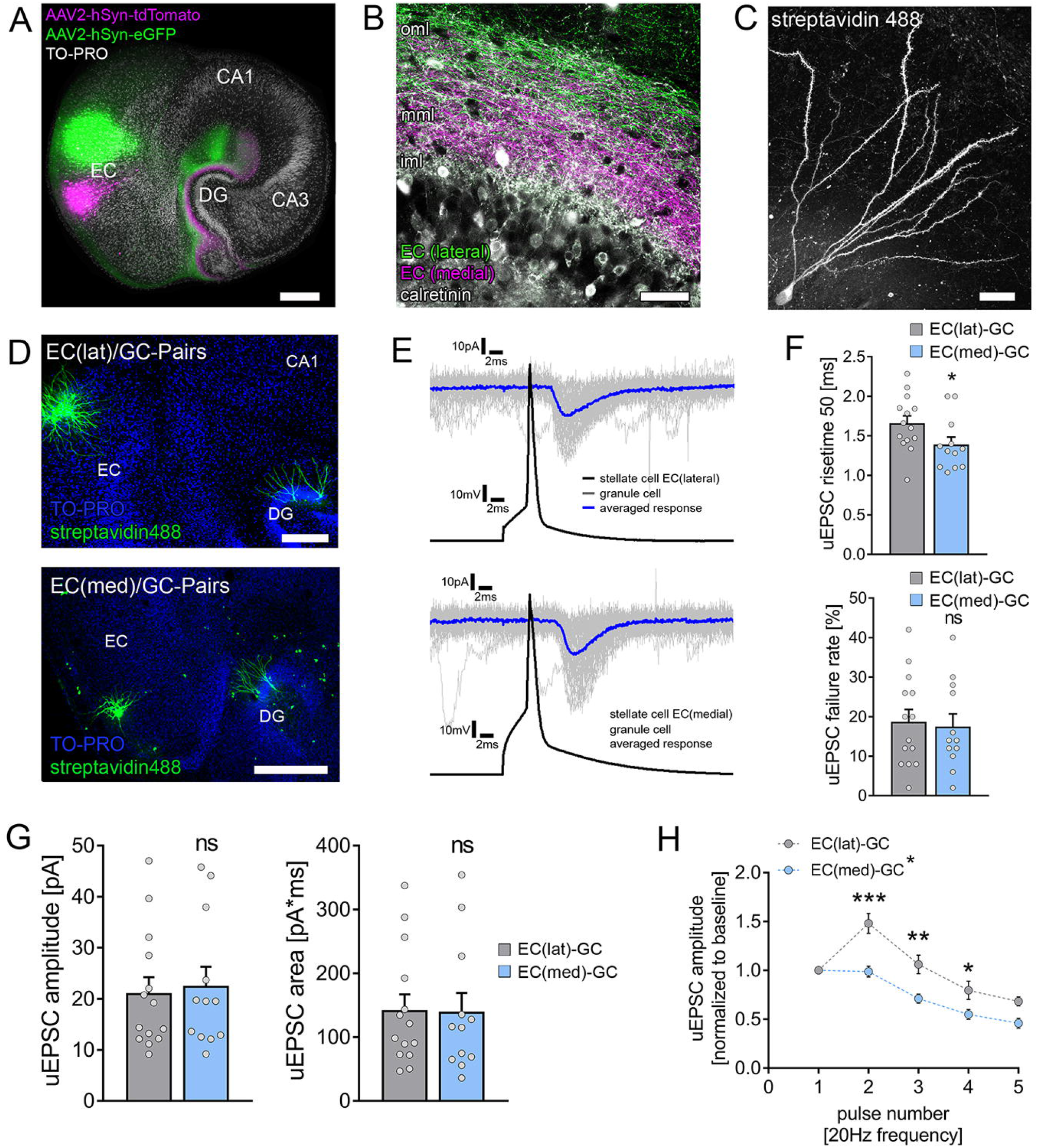
Functional characterization of laminar innervation of dentate granule cells in organotypic entorhino-hippocampal slice cultures. (A) Overview of a mouse organotypic entorhino-hippocampal tissue culture stained with TO-PRO nuclear stain (white) to visualize cytoarchitecture. The entorhino-hippocampal projection is visualized by a perforant path tracing using AAV-mediated expression of tdTomato (medial part) and eGFP (lateral part) in the entorhinal cortex. Scale bar = 300 μm. DG, dentate gyrus; EC, entorhinal cortex. (B) The dentate gyrus at higher magnification demonstrating its laminar innervation. eGFP labeled axonal projections from the lateral part of the entorhinal cortex reach the outer molecular layer (oml), while tdTomato labeled fibers project to the middle molecular layer (mml). The inner molecular layer (iml) that contains hilar mossy cell axons (visualized by calretinin immunostaining) is not innervated by entorhino-hippocampal projections. Scale bar = 100 μm. (C) Posthoc streptavidin-stained dentate granule cell. Scale bar = 20 μm. (D) Posthoc-staining of paired recordings from layer 2 stellate cells in either the lateral (upper panel) or the medial part (lower panel) of the entorhinal cortex and dentate granule cells in the suprapyramidal blade of the dentate gyrus. TO-PRO^®^ nuclear stain was used to visualize cytoarchitecture. Scale bar = 200 μm (upper panel) and 300 μm (lower panel). DG, dentate gyrus; EC, entorhinal cortex. (E) Action potentials were induced in EC stellate cells (50 action potentials, black trace) and postsynaptic responses were recorded in dentate granule cells (gray traces, single sweeps; blue trace, averaged response). (F) Synaptic failure rate was not significantly different between paired recordings from the lateral and the medial part of the perforant path. Nevertheless, monosynaptic connections between the medial layer 2 stellate cells and dentate granule cells had significantly faster risetimes (n_EC(lat)-GC_ = 14 pairs in 5 independent experiments; n_EC(med)-GC_ = 12 pairs in 5 independent experiments; Mann-Whitney test, U = 41). (G) No differences in uEPSC amplitude or area were found between connected neuronal pairs of the lateral and the medial perforant path (Mann-Whitney test). (H) Short-term plasticity was assessed by the induction of 5 action potentials at 20 Hz frequency in presynaptic stellate cells. EC(lat)-GC pairs displayed a synaptic facilitation while EC(med)-GC showed neither synaptic facilitation nor depression in response to the second presynaptic action potential. Repetitive stimulation of both pathways resulted in progressive synaptic depletion. Thus, significant differences between the individual pathways upon repetitive presynaptic stimulation at the synaptic level were identified (RM two-way ANOVA with Sidak’s multiple comparisons test). Individual data points are indicated by gray dots. Values represent mean ± s.e.m. (*, p < 0.05; **, p < 0.01; ***, p < 0.001; ns, non-significant difference).

### Paired whole-cell patch clamp recordings reveal lamina-specific differences in short-term plasticity of perforant path synapses

To characterize the entorhino-hippocampal projections at the level of individual connected pairs of neurons, layer 2 stellate cells in the lateral or the medial part of the entorhinal cortex and granule cells in the suprapyramidal blade of the dentate gyrus were patched simultaneously (Figure 1C, D). Solitary action potentials were induced in the presynaptic stellate cell while recording excitatory postsynaptic responses from the soma of individual granule cells (Figure 1E). Neurons were considered to be connected if at least 5% of the presynaptic action potentials induced time-locked unitary excitatory postsynaptic currents (uEPSC) in the dentate granule cell (c.f. (Vlachos et al., 2012c)). Both, neurons in the lateral and the medial part of the entorhinal cortex formed direct synaptic connections onto dendrites of dentate granule cells. Our analysis revealed that the uEPSC risetime was significantly lower in the recorded pairs of medial stellate cells and dentate granule cells as compared to lateral stellate cells (Figure 1F). In line with previous observations that report a positive correlation between risetime and the synaptic distance from the soma (Lenz et al., 2019; Sjostrom and Hausser, 2006; Vlachos et al., 2012b), this functional result supported our structural analysis, i.e., proper lamination of excitatory inputs onto dentate granule cells (c.f., Figure 1A, B).

The synaptic failure rate, which reflects the amount of presynaptic action potentials that fail to elicit postsynaptic responses, was not different between the medial and the lateral perforant path synapses (Figure 1F). Moreover, the uEPSC amplitude and the area were not significantly different between the two pathways (Figure 1G), suggesting that medial and lateral perforant path synapses equally affect somatic membrane potentials. However, the assessment of short-term plasticity through the induction of five consecutive action potentials (at 20 Hz) in the presynaptic cells revealed that the lateral perforant path expressed robust short-term facilitation which was not seen in the medial perforant path synapses. We conclude that the ability to express this form of short-term plasticity distinguishes perforant path synapses in the oml and mml of the dentate gyrus (Figure 1H).

### Hilar mossy cell fibers in the inner molecular layer form highly reliable synapses onto dentate granule cells

In order to assess synaptic properties of synapses close to the soma of granule cells, paired recordings between hilar mossy cells and dentate granule cells were carried out (Figure S1A). Monosynaptically connected pairs between hilar mossy cells and dentate granule cells showed high uEPSC amplitudes, fast risetimes, and low synaptic failure rates (Figure S1B, C). Notably, short-term plasticity experiments revealed a strong depression and fast depletion behavior for mossy cell-to-dentate granule cell synapses (Figure S1D). We conclude that short-term synaptic plasticity switches from facilitation on distal dendrites to depression on proximal dendrites of dentate granule cells.

### Computational modeling of excitatory synaptic inputs indicates a comparable depolarizing impact of the medial and the lateral perforant path on dentate granule cells

Previous work demonstrated that dentate granule cell dendrites strongly attenuate synaptic signals (Krueppel et al., 2011). However, we did not observe differences in uEPSC amplitudes originating from synapses in the oml and mml (c.f., Figure 1G). This finding was further explored by computing the dependence of EPSC amplitude to distance from soma using compartmental modeling of mouse dentate granule cells. To determine the distribution of simulated EPSCs with increasing distance from the granule cell layer, identical single synaptic inputs were activated at different locations along a path between the soma and a distal end of the dendrite. Corresponding EPSC responses were detected at the soma and the simulated cells were voltage clamped at −70 mV – similar to our experiments (Figure S2A). We found a gradual reduction of somatically recorded EPSCs along the dendritic path. This dendritic attenuation stabilized between 100 μm and 200 μm distance from the soma, which corresponds to the outer two thirds of the molecular layer, i.e., mml and oml. Hence, perforant path synapses of comparable synaptic strength in the mml and oml are expected to equally depolarize the soma of dentate granule cells. Moreover, somatic EPSCs were simulated for the simultaneous activation of excitatory synapses with equal distances from the soma on all granule cell dendrites. A plateau in somatic EPSC responses was detected that correlated with the layer-specific dendritic complexity (Figure S2B). Again, these data suggested that the magnitude of somatic EPSC responses from dendritic synaptic activation remains stable over a long range of middle to long distances from the soma.

### APP regulates excitatory synaptic transmission at medial perforant path synapses

The amyloid precursor protein and its cleavage products have been recognized as regulators of synaptic transmission (Muller et al., 2017). Beyond its role in pathological processes, in which deficits at the medial perforant path have been reported, recent studies suggest a physiological role of APP and its fragments. Thus, we here addressed the question of whether APP-deficiency affects synaptic transmission between monosynaptically connected medial layer 2 stellate cells and mature dentate granule cells (Figure 2A). First, APP-deficiency was accompanied by a decrease in uEPSC failure rate, i.e., an increased synaptic reliability (Figure 2B). Moreover, the uEPSC amplitude and area were significantly increased in the absence of APP (Figure 2C). The short-term plasticity experiments confirmed increased uEPSC amplitudes and revealed a synaptic depression of medial perforant path synapses in the APP-deficient dentate gyrus (Figure 2D). We conclude that APP is involved in regulating the reliability and strength of medial perforant path synapses.

**Figure 2:**
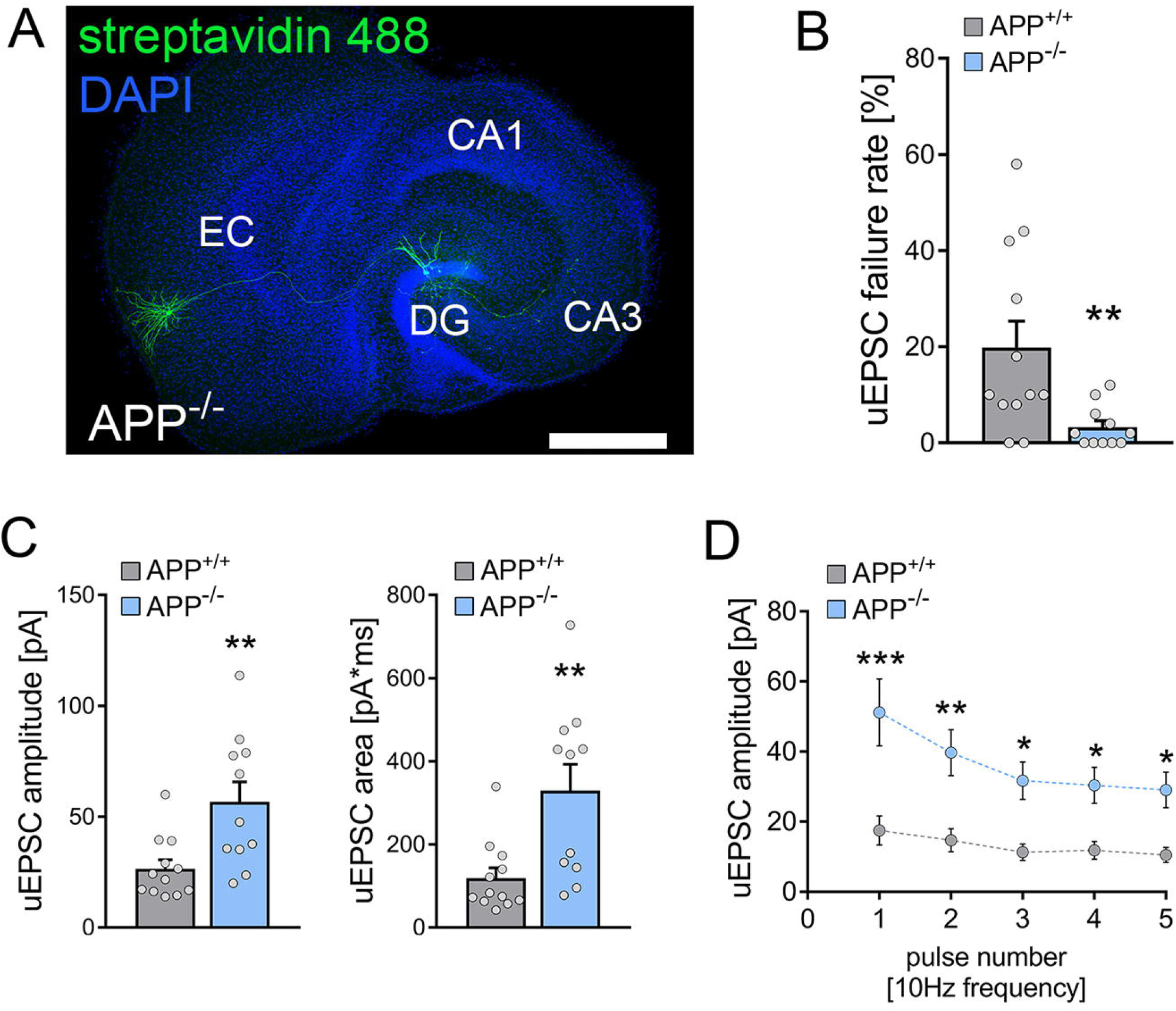
The amyloid precursor protein (APP) regulates excitatory synaptic transmission at medial perforant path synapses. (A) Posthoc-staining of paired recordings from layer 2 stellate cells in the medial part of the entorhinal cortex and dentate granule cells in the suprapyramidal blade of the dentate gyrus in APP-deficient (APP^−/−^) cultures. DAPI nuclear stain was used to visualize cytoarchitecture. Cultures from APP-deficient animals have no overt morphological aberration of the perforant path. Scale bar = 500 μm. DG, dentate gyrus; EC, entorhinal cortex. (B) The uEPSC failure rate was significantly reduced in APP^−/−^ cultures compared to wildtype (APP^+/+^) cultures (n_wildtype_ = 12 pairs in 5 independent experiments, n_APP-deficient_ = 11 pairs in 5 independent experiments; Mann-Whitney test, U = 25.5). (C) Both uEPSC amplitude and area were significantly increased in APP^−/−^ cultures (Mann-Whitney test, U_uEPSC amplitude_ = 23, U_uEPSC area_ = 19). (D) Assessment of short-term plasticity by the application of five presynaptic action potential at 10 Hz frequency confirmed the increase in uEPSC amplitude with synaptic depletion upon repetitive stimulation in APP^−/−^ cultures (RM two-way ANOVA with Sidak’s multiple comparisons test). Individual data points are indicated by gray dots. Values represent mean ± s.e.m. (*, p < 0.05; **, p < 0.01; ***, p < 0.001).

### Transcriptomic changes in the APP-deficient dentate gyrus

To further characterize the effects of APP-deficiency in the dentate gyrus of entorhino-hippocampal tissue culture, we performed a region specific transcriptome analysis of dentate gyrus samples isolated from wildtype and APP-deficient tissue cultures (Figure 3A-C). APP-deficiency was associated with complex changes in the gene expression profile within the dentate gyrus (Figure 3A), where numerous differentially expressed genes were detected (UP: 911 genes, DOWN: 1206 genes, UNCHANGED: 9039 genes; Table S1). A gene set enrichment analysis for cellular compartment gene ontologies revealed that the differential gene expression can be attributed to distinct clusters, e.g., mitochondria, endoplasmic reticulum, and ribosomes (Figure 3B). Notably, a cluster for differential gene expression was also found for synapse related genes (Figure 3B, bold), where differential expression of genes related to postsynapses, presynapses, and synaptic vesicles could be identified (Figure 3C). Thus, APP-deficiency leads to complex changes in neural gene expression in the dentate gyrus that might contribute to alterations in excitatory synaptic transmission and plasticity.

**Figure 3:**
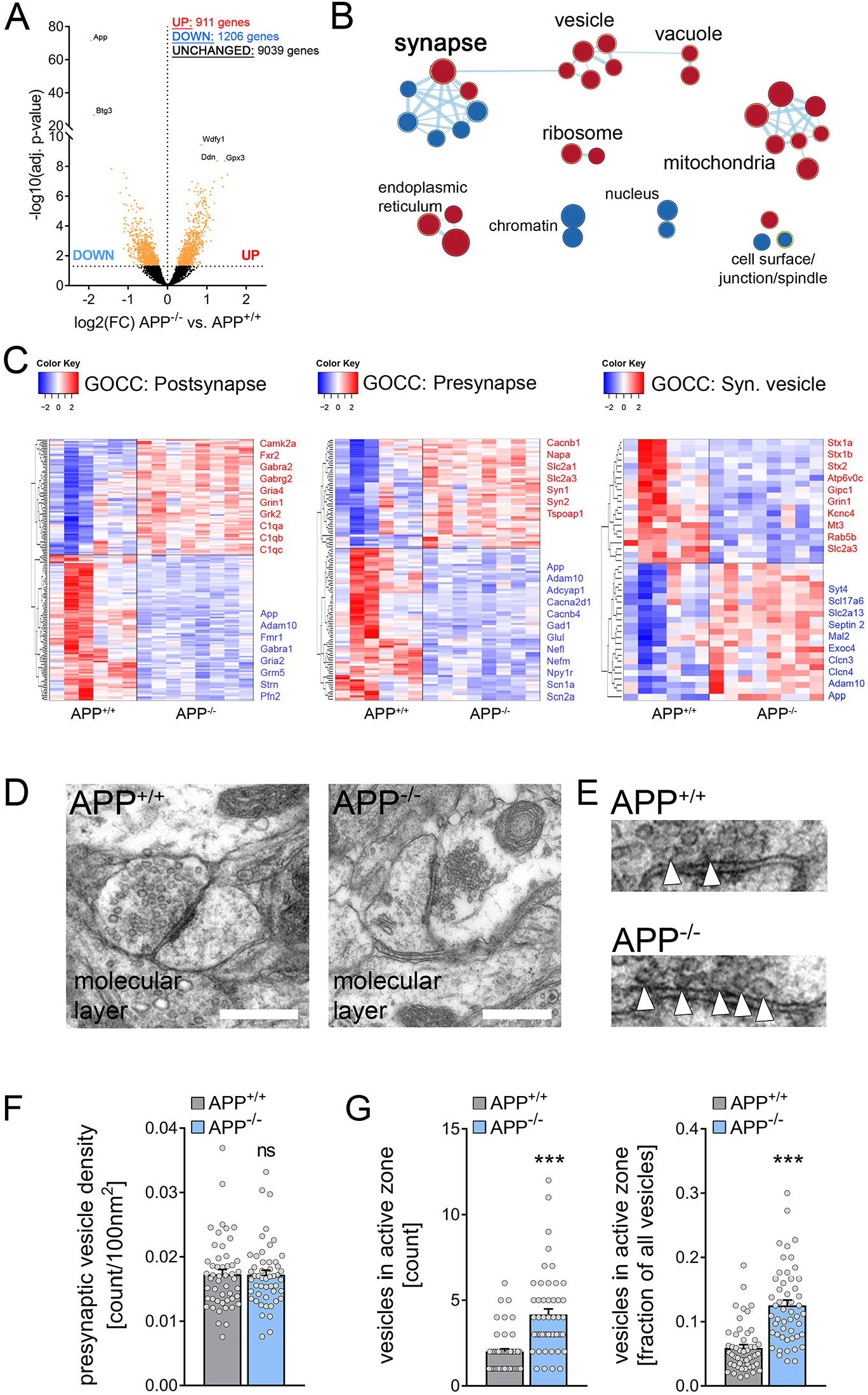
APP-deficiency modulates the dentate gyrus transcriptome and is accompanied by ultrastructural changes in perforant path presynaptic terminals. (A) Volcano-plot from differential gene expression analysis using DeSeq2 comparing gene expression levels in the dentate gyrus of wildtype (APP^+/+^) and APP-deficient cultures (APP^−/−^). Differentially expressed genes that passed the read number threshold (BaseMean > 150 normalized reads in DeSeq2 output) were indicated with colored dots (n_APP+/+_ = 6 samples, n_APP-/-_ = 8 samples; significance threshold at p(adj.) = 0.05). (B) Clusters of differentially expressed genes were identified by gene set enrichment analysis (p < 0.05; FDR threshold 0.1). Cluster interaction is indicated by connection line thickness and cluster size corresponds to the circle size. Colors indicate positive (red) and negative (blue) correlations of differential gene expression. Clusters were grouped manually for comprehensive illustration. (C) Heatmaps for differentially expressed genes dedicated to cellular compartment gene ontologies (GOCC; postsynapse, presynapse, synaptic vesicle). Selected genes are highlighted next to the respective heatmap. (D) Ultrastructural features of presynaptic perforant path terminals were assessed in electron micrographs of the molecular layer in wildtype (APP^+/+^) and APP deficient (APP^−/−^) cultures. Scale bars = 500 nm. (E) Synaptic vesicles were counted in presynaptic buttons. Moreover the number of vesicles attached to the presynaptic active zone was assessed (white arrowheads). (F) No difference in presynaptic vesicle density between APP^+/+^ and APP^−/−^ preparations was detected (n = 50 presynaptic terminals in each group in 3 independent experiments; Mann-Whitney test). (G) Both the total number of docked vesicles in the presynaptic active zone and the fraction of docked vesicles from the total number of vesicles were increased in APP-deficient tissue preparations (Mann-Whitney test, U_count_ = 442.5, U_fraction_ = 386). Individual data points are indicated by gray dots. Values represent mean ± s.e.m. (***, p < 0.001; ns, non-significant difference).

### APP-deficiency is accompanied by ultrastructural changes of presynaptic sites at perforant path synapses

To test whether the APP-deficiency was further accompanied by structural changes at pre- and postsynaptic compartments that might correspond to the functional and transcriptomic changes, transmission electron microscopy of individual synapses in the molecular layer was employed (Figure 3D). Of note, previous studies did not show any significant postsynaptic changes in APP-deficient tissue cultures (Galanis et al., 2021). The analysis of presynaptic features revealed that the presynaptic vesicle density is unaltered between the groups (Figure 3E, F). In contrast, both the number and the fraction of vesicles in the presynaptic active zone were significantly increased in APP-deficient tissue cultures (Figure 3G). Therefore, we conclude that APP-deficiency is accompanied by an increased amount and fraction of ready-to-release vesicles at perforant path synapses onto dentate granule cells.

### Presynaptic but not postsynaptic APP modulates excitatory neurotransmission at medial perforant path synapses

Finally, we addressed the impact of pre- and postsynaptic APP expression on excitatory synaptic transmission at medial perforant synapses. In these experiments, tissue cultures from APP^flox/flox^ x Ai14 Cre reporter mice were used and recombination of the floxed APP gene and expression of Ai14 Cre-dependent tdTomato was achieved through the local viral injection of AAV-hSyn-Cre-GFP. Postsynaptic deletion of APP was realized by viral infections in the dentate gyrus (Figure 4A, left panel), and presynaptic APP deletion by Cre-GFP expression in the medial part of the entorhinal cortex in a different set of tissue cultures (Figure 4A, right panel). Once more, individual connected pairs of medial layer 2 stellate cells and dentate granule cells were patched and cellular recombination was confirmed by the somatic tdTomato fluorescence signal during the recordings (Figure 4A, B). The presynaptic APP-deficiency was accompanied by a significant reduction in synaptic failure rates (Figure 4C) and a robust increase in both the amplitude and area of successfully evoked uEPSC was observed (Figure D). Conversely, postsynaptic recombination did not significantly affect synaptic transmission although a trend towards a decrease in uEPSC failure rate should be mentioned (p = 0.07; Figure 4C, D). Notably, short-term plasticity experiments demonstrated that both pre- and postsynaptic recombination were associated with synaptic depression and increased synaptic depletion compared to wildtype tissue cultures, while only presynaptic APP-deficiency resulted in higher uEPSC amplitudes (Figure 4E). Thus, we conclude that excitatory neurotransmission at medial perforant path synapses is effectively restrained by presynaptic but not postsynaptic APP.

**Figure 4:**
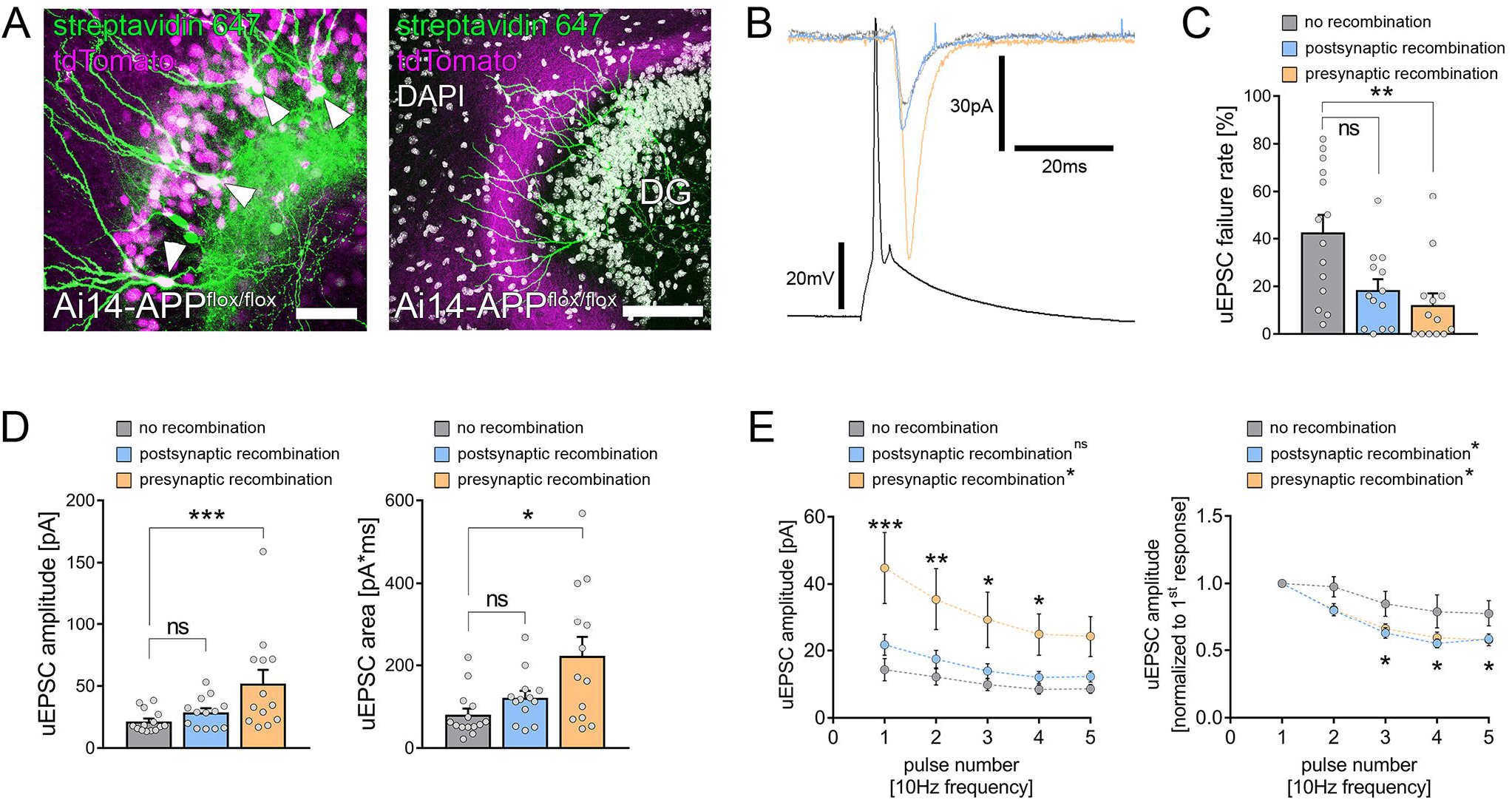
Presynaptic APP restrains excitatory neurotransmission at medial perforant path synapses. (A) To achieve compartment specific deletion of APP, tissue cultures from Ai14-APP^flox/flox^ animals were prepared and region specific recombination was achieved by local injection of a Cre-expressing virus in either the dentate gyrus (left panel) or the medial part of the entorhinal cortex (right panel). Granule cells (arrowheads, posthoc-staining with streptavidin 647 of cells that were filled with biocytin during the recording) were patched and recombination was evaluated during the patch-clamp experiment by the tdTomato signal. Scale bars = 50 μm (left panel), 100 μm (right panel). (B) Electrophysiological properties were assessed in monosynaptic connected pairs of neurons with no recombination (gray line), postsynaptic (blue trace) or presynaptic (orange trace) recombination. (C) The uEPSC failure rate was significantly decreased in connected pairs with presynaptic APP-deficiency, while postsynaptic APP-deficiency resulted in a non-significant trend toward increased synaptic reliability (n_no recombination_ = 14 pairs in 5 independent experiments; n_postsynaptic recombination_ = 13 pairs in 5 independent experiments; n_presynaptic recombination_ = 13 pairs in 5 independent experiments; Kruskal-Wallis test followed by Dunn’s multiple comparisons test). (D) Presynaptic, but not postsynaptic recombination was accompanied by an increase in both uEPSC amplitude and area when compared with non-recombined pairs of neurons (Kruskal-Wallis test followed by Dunn’s multiple comparisons test). (E) Assessment of short-term plasticity confirmed the increase in uEPSC amplitude upon presynaptic but not postsynaptic recombination. Moreover, normalized analysis revealed a more prominent synaptic depletion after both pre- and postsynaptic recombination (RM two-way ANOVA with Sidak’s multiple comparisons test). Individual data points are indicated by gray dots. Values represent mean ± s.e.m. (*, p < 0.05; **, p < 0.01; ***, p < 0.001; ns, non-significant difference).

## DISCUSSION

The eminent role of the perforant path in normal brain function and pathological processes, such as Alzheimer’s disease, has been extensively studied (Hainmueller and Bartos, 2018; Hyman et al., 1986; Robinson et al., 2014; Witter, 2007). Nevertheless, a characterization at the level of individual connected cells, i.e. layer 2 stellate cells and dentate granule cells, remained challenging due to the complex three dimensional organization of this pathway. Organotypic tissue cultures of the entorhino-hippocampal complex provide the opportunity to investigate cortico-hippocampal projections in a laminated and steady-state environment characterized by *in vivo*-like ultrastructural features of excitatory synapses (Lenz et al., 2021; Maus et al., 2020). Our results demonstrate that unitary synaptic events at lateral and medial perforant path synapses are not distinct with respect to their depolarizing effects on the soma of dentate granule cells. However, trains of action potentials revealed lamina-specific differences in short-term plasticity. Since these features are evident at the level of individually connected neurons, we conclude that differences in short-term plasticity at distinct excitatory inputs onto dentate granule cells are related to lamina specific synaptic properties rather than being a network phenomenon. Moreover, the properties of medial perforant path synapses depended on the presence of APP, since APP-deficiency significantly enhanced excitatory synaptic transmission by decreasing synaptic failure rates and increasing the amplitudes and areas of uEPSCs. Consistently, increased numbers of docked vesicles at presynaptic active zones were observed. Finally, we provide evidence that presynaptic but not postsynaptic APP-deficiency significantly enhanced excitatory synaptic transmission suggesting a compartment specific regulatory role of APP.

The amyloid precursor protein (APP) is widely expressed in the central nervous system (Bergstrom et al., 2016; Del Turco et al., 2016). Beyond the link to pathological conditions such as Alzheimer’s Disease (AD) its physiological role in synaptic transmission has been recognized (Jedlicka et al., 2012; Muller et al., 2017). A notable feature of the full-length protein is its proteolytic processing by so-called secretases (e.g. Bace-1 and Adam10) that results in fragments with distinct functions in regulating neural circuits (Jimenez et al., 2011; Nunan and Small, 2000; Vassar et al., 1999). While Aβ-fragments were mostly linked to pathological conditions (Opazo et al., 2018), neuroprotective and synaptic plasticity enhancing properties have been attributed to soluble APPα (Bold et al., 2022; Fol et al., 2016; Jimenez et al., 2011; Mockett et al., 2017; Richter et al., 2018; Tan et al., 2018). Moreover, recent advances demonstrated that APP and its fragments are regulators of homeostatic synaptic plasticity which aims at keeping neuronal activity in a dynamic range through synaptic adaptations (Galanis et al., 2021). Since APP has an impact on the homeostatic regulation of synaptic features, APP-deficiency might lead to maladaptive plasticity and excitatory synaptic enhancement at medial perforant path synapses (Maggio and Vlachos, 2014).

As revealed by previous studies, APP can be found at the presynaptic active zone and has been suggested to have dedicated roles in the synaptic vesicle cycle (Lassek et al., 2013; Lassek et al., 2016; Wang et al., 2005; Yang et al., 2005). Moreover, APP and its cleavage products can affect hippocampal networks (Harris et al., 2010). The release of Aβ from perforant path terminals has been recognized to affect synaptic integrity (Lazarov et al., 2002) that can depress excitatory synapses (He et al., 2019; Kamenetz et al., 2003). In this context, presynaptic calcium channels and vesicle recycling represent targets for Aβ which mediates a suppression of spontaneous synaptic activity (Kelly and Ferreira, 2007; Kelly et al., 2005; Nimmrich et al., 2008). In line with these findings, we here provide evidence that presynaptic APP – which might promote localized actions of Aβ at presynaptic sites – is essential to restrain excitatory synaptic transmission at medial perforant path synapses on dentate granule cells. These findings support recent evidence on the role of presynapses in the pathogenesis of Alzheimer’s disease (Barthet and Mulle, 2020; Jorda-Siquier et al., 2022). In support of a restraining effect of APP and its proteolytic cleavage products, we here show at the level of connected pairs of neurons that presynaptic APP controls the efficacy and strength of perforant path synapses. Indeed, ultrastructural analysis confirmed our functional data by demonstrating increased numbers of docked vesicles in the active zone of synapses. In contrast, other studies have revealed decreases in presynaptic vesicle abundance at neuromuscular synapses in genetic models targeting several APP-related genes (e.g. double mutants for APLP2 and APP; (Wang et al., 2005; Yang et al., 2005)) indicating a complex regulation of presynaptic function by members of the APP protein family. Whether APP functions as an uncleaved transmembrane protein or via amyloidogenic or anti-amyloidogenic processing products warrants further investigation.

Presynaptic APP levels are regulated through anterograde axonal transport. Several studies have shown that anterograde axonal transport is affected early in neurodegenerative diseases which can promote amyloidogenesis (Bera et al., 2020; Chaves et al., 2021; Tang, 2009). Moreover, APP levels in the presynaptic compartment decrease upon disturbances in axonal transport during neurodegeneration (Morotz et al., 2019). Based on our results, the reduction in APP levels in presynaptic terminals of the perforant path might account for a synaptic imbalance and maladaptation through excitatory synaptic enhancement. Notably, recent studies report that alterations in presynaptic APP levels can be correlated to changes in postsynaptic excitatory synapse morphologies (Bruyere et al., 2020). In line with these findings, hyperexcitability of neural networks is commonly observed in animal models and human subjects with Alzheimer’s disease (Ghatak et al., 2019; Voskobiynyk et al., 2020; Zott et al., 2019). Thus, it is interesting to speculate that defects in the axonal transport of APP during Alzheimer’s disease with a subsequent decrease in presynaptic APP levels might contribute to alterations in information processing (Haytural et al., 2020). In line with this hypothesis, previous studies reported alterations in network activity upon APP-deficiency (Zhang et al., 2016). Nevertheless, APP-deficient mice are behaviorally normal at a young age which suggest distinct mechanisms that compensate for the lack of APP (Muller et al., 2017).

Information processing in the dentate gyrus is considered as a crucial step during memory acquisition. In this regard, the pattern separation properties of dentate granule cells have been linked to performances in cognitive tasks (Bekinschtein et al., 2013; Frank et al., 2020; Leutgeb et al., 2007). The role of short-term-plasticity and synaptic strength at perforant path synapses for the computations of the dentate gyrus remain not well understood. Our results demonstrate that compartmentalized APP-deficiency significantly enhanced excitatory neurotransmission at medial perforant path synapses onto dentate granule cells. We hypothesize that these alterations in excitatory perforant path neurotransmission might significantly interfere with the pattern separation abilities of the dentate gyrus network. It is interesting to speculate that behavioral deficits in Alzheimer’s disease might result from presynaptically driven altered information processing in the dentate gyrus (Barthet and Mulle, 2020). Thus, we conclude that APP-deficiency at presynaptic terminals – which might be caused by genetic loss or defects in axonal transport – affect hippocampal information processing and memory formation processes.

## Acknowledgements

We thank Dr. R. Jude Samulski and the UNC Vector Core for providing adeno-associated viruses (AAV2-hSyn-GFP-Cre). We thank Simone Zenker and Sigrun Nestel for skilful technical assistance. The Galaxy server used for some calculations is in part funded by Collaborative Research Centre 992 Medical Epigenetics (DFG grant SFB 992/1 2012), and the German Federal Ministry of Education and Research (BMBF grants 031 A538A/A538C RBC, 031L0101B/031L0101C de.NBI-epi, 031L0106 de.STAIR (de.NBI)). This work was supported by Else Kröner-Fresenius-Stiftung (EKFS_#2019_A94 to M.L.), Deutsche Forschungsgemeinschaft (FOR 1332 to U.M., T.D., and A.V., and CRC/TRR 167 Project-ID 259373024 B14 to A.V.) and the German Federal Ministry of Education and Research (OGEAM-2 16LW0161K to U.M. and to T.D.).

## Author contributions

Author contributions have been assigned according to CRediT taxonomy. ML: Conceptualization, Methodology, Validation, Formal Analysis, Investigation, Writing-original draft preparation, Visualization, Project administration, Funding acquisition. AE: Investigation, Formal Analysis. PK: Investigation, Formal Analysis. CG: Formal Analysis. DK: Formal Analysis. PJ: Investigation, Formal Analysis. UM: Resources. TD: Resources. AV: Conceptualization, Methodology, Resources, Writing-original draft preparation, Supervision, Project administration. All authors contributed to critical review and comments to the final version of the manuscript before submission.

