## Supplementary Information for "The amyloid precursor protein regulates synaptic transmission at medial perforant path synapses"

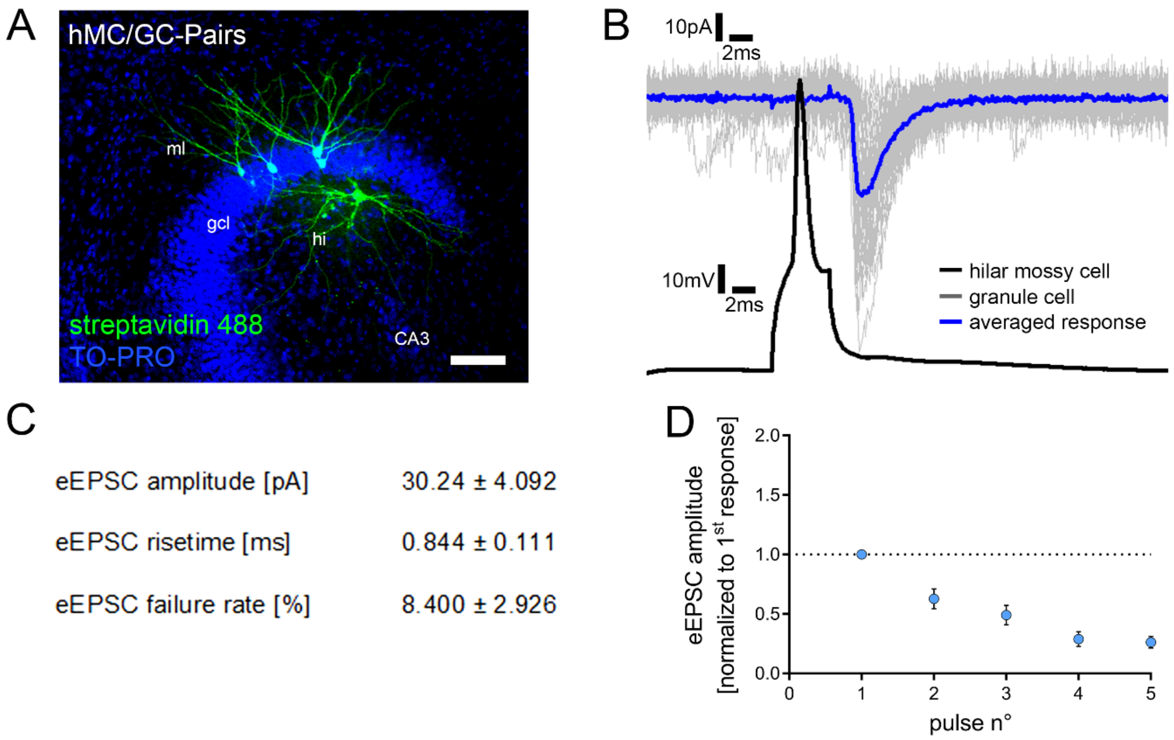


Figure S1

**Figure S1: Single-cell characterization of the hilar mossy cell – dentate granule cell projection.**

(A) Posthoc-staining of paired recordings from hilar mossy cells (hMC) and dentate granule cells in the suprapyramidal blade of the dentate gyrus. TO-PRO nuclear stain was used to visualize cytoarchitecture. Hilar mossy cells were identified during the experiment by their morphological and electrophysiological features. Scale bar = 100 µm. gcl, granule cell layer; ml, molecular layer; hi, hilar region.

(B) Action potentials were induced in hilar mossy cells (50 action potentials, black trace) and postsynaptic responses were recorded in dentate granule cells (gray traces, single sweeps; blue trace, averaged response). Connected pairs showed highly reliable inward currents upon presynaptic stimulation.

(C) Summary table for single-cell characterization of excitatory inputs onto dentate granule cells, originating from hilar mossy cells (5 pairs in 21 cultures).

(D) Short-term-plasticity experiments revealed a depressive and depletive behavior of this connection following repetitive stimulation.

Values represent mean ± standard error of the mean.

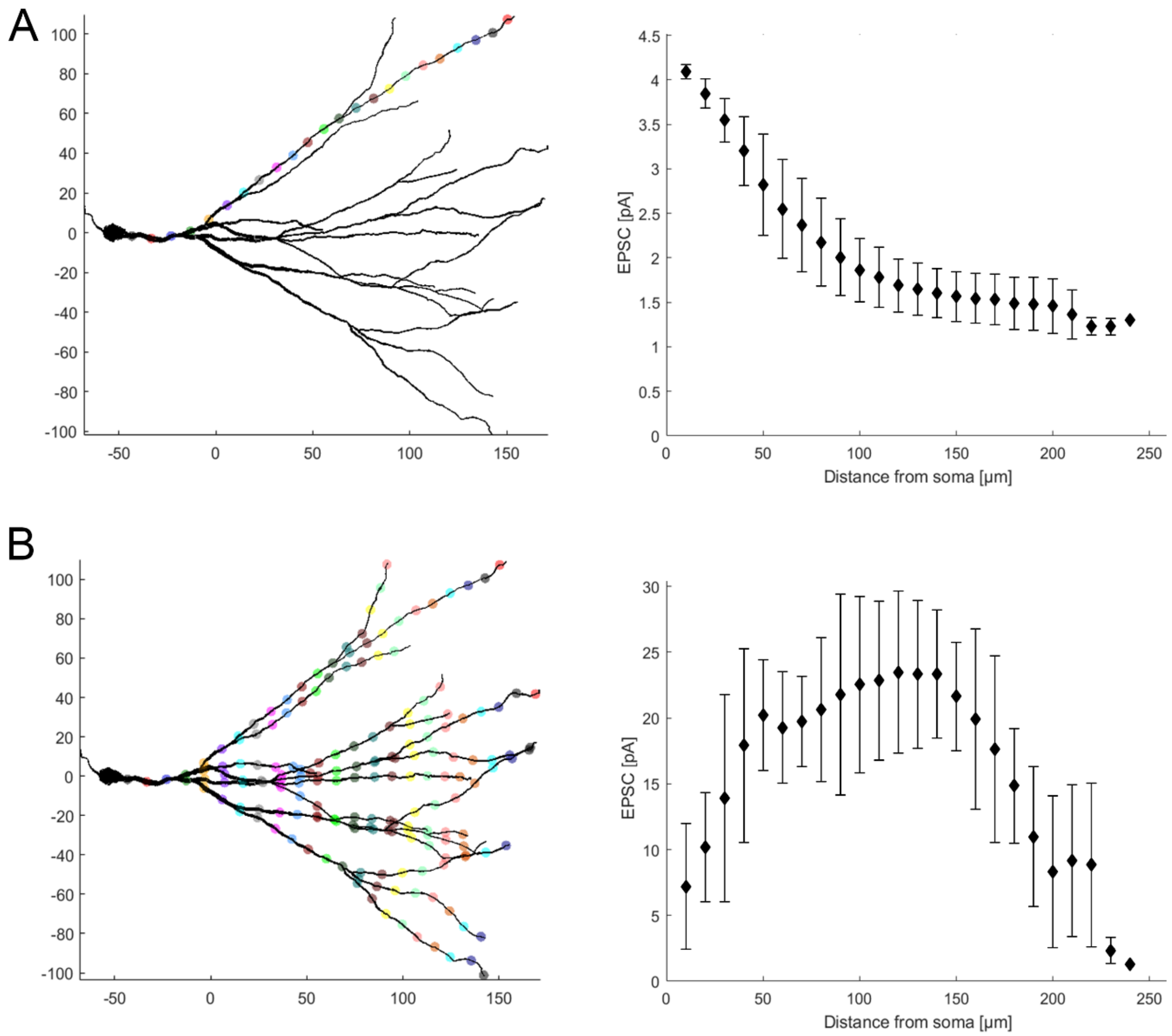


Figure S2

**Figure S2: Computational modeling of somatic currents elicited by synaptic activation on dentate granule cell dendrites.**

(A) Modeling of somatic currents elicited by a single synaptic activation along a specific granule cell dendrite. Colored dots indicate the location of individual synaptic activation. The right panel shows the *in silico* recorded somatic EPSC depending on the synaptic distance from the granule cell soma.

(B) Modeling of somatic currents elicited by the simultaneous activation of synapses at granule cell dendrites at distinct distances from the soma. Simultaneously activated synapses are depicted in the same colors. The right panel shows the *in silico* recorded somatic EPSC depending on the synaptic distance from the granule cell soma.

Values represent mean ± standard error of the mean.
